# A mechanistically interpretable model of the retinal neural code for natural scenes with multiscale adaptive dynamics

**DOI:** 10.1101/2021.12.18.473320

**Authors:** Xuehao Ding, Dongsoo Lee, Satchel Grant, Heike Stein, Lane McIntosh, Niru Maheswaranathan, Stephen Baccus

## Abstract

The visual system processes stimuli over a wide range of spatiotemporal scales, with individual neurons receiving input from tens of thousands of neurons whose dynamics range from milliseconds to tens of seconds. This poses a challenge to create models that both accurately capture visual computations and are mechanistically interpretable. Here we present a model of salamander retinal ganglion cell spiking responses recorded with a multielectrode array that captures natural scene responses and slow adaptive dynamics. The model consists of a three-layer convolutional neural network (CNN) modified to include local recurrent synaptic dynamics taken from a linear-nonlinear-kinetic (LNK) model [1]. We presented alternating natural scenes and uniform field white noise stimuli designed to engage slow contrast adaptation. To overcome difficulties fitting slow and fast dynamics together, we first optimized all fast spatiotemporal parameters, then separately optimized recurrent slow synaptic parameters. The resulting full model reproduces a wide range of retinal computations and is mechanistically interpretable, having internal units that correspond to retinal interneurons with biophysically modeled synapses. This model allows us to study the contribution of model units to any retinal computation, and examine how long-term adaptation changes the retinal neural code for natural scenes through selective adaptation of retinal pathways.

## I. Introduction

The processing of visual information in the nervous system includes a broad set of computations, with timescales that range from milliseconds to tens of seconds. On a fast timescale, the precise timing of fast neural dynamics with a short latency is essential to encode immediate properties of the stimulus like the trajectory of a moving object. Over longer timescales, adaptive properties produced by short-term plasticity measure the statistics of the stimulus, allowing cells to use their dynamic range efficiently and improve information transmission [2]. One well-known example of computations that span multiple timescales is contrast adaptation [3], [4], whereby an increase in contrast triggers both fast and slow adjustments in sensitivity and response dynamics. To understand these neural computations, dynamics and their mechanisms in a unified framework, it is important to create a quantitative multi-timescale neural encoding model with components that can be mapped to neural mechanisms.

The challenge posed by modeling this broad range of neural dynamics has thus far been addressed with separate types of models. As shown in [5], [6], CNN models of the retina are capable of predicting neural responses accurately for stimuli of nontrivial spatiotemporal structures as well as capturing many phenomena within the retinal integration timescale. Moreover, they are interpretable in that internal model units are highly correlated with the membrane potential of interneurons. However, constrained by the feed-forward nature of these models, long-timescale phenomena induced by synaptic plasticity are beyond their predicting capabilities. In contrast, the LNK model [1] successfully captures contrast adaptation over multiple timescales using a first-order continuous Markov model of the type used to describe chemical reactions, ion channels, and synapses, but this model has been used only for a spatially uniform stimulus. Consequently, it is unclear what role such kinetics can play in a more realistic model setting and how they interact with spatial information processing in the retina.

Here we present a recurrent model of salamander retinal ganglion cells consisting of two convolutional layers and local recurrent synaptic dynamics taken from the LNK model [1]. These synaptic units are interpretable in that they correspond to biophysical mechanisms of synaptic vesicle release. We designed a novel protocol to train both the fast spatiotemporal parameters and slow synaptic parameters of the model separately using natural scenes and spatially uniform white noise data sets. The resulting trained model is capable of predicting the firing rates of retinal ganglion cells accurately and capturing contrast adaptation as well as other visual computations that our previous CNN models also capture [6]. Furthermore, our results suggest a general approach to fitting a single model with interpretable recurrent synaptic units to capture neural dynamics over multiple timescales.

## II. Methods

### A. Visual stimuli

A video monitor projected visual stimuli at 30 Hz controlled by Matlab (Mathworks), using Psychophysics Toolbox [7], [8]. Stimuli had a constant mean intensity of 8.3*mW/m*^2^. Images were presented in a 50×50 grid with a square size of 55*µm*.

We presented alternating natural scenes and spatially uniform white noise stimuli. The natural scene stimulus was a sequence of jittered natural images sampled from a natural image database [9]. The stimulus intensity of the spatially uniform white noise was drawn every 33 ms from Gaussian distributions whose contrast (standard deviation / mean intensity) changed every 20 seconds. The contrast of each 20*s* block was drawn uniformly from the interval [0.05, 0.35].

### B. Electrophysiology

The responses of tiger salamander retinal ganglion cells were recorded using a 60 channel multielectrode array. Further experimental details are described in detail elsewhere [10].

### C. Data preparation

Spiking responses were binned using 10 ms bins and smoothed using a 10 ms Gaussian filter. The training dataset of 95 minutes in total was divided according to a 90%*/*10% train/validation split. The test dataset consisted of averaged repeated trials to novel stimuli of 60 seconds natural scene and 220 seconds spatially uniform white noise.

### D. Model architecture

As shown in Fig. 1, the stimulus was convolved with eight spatiotemporal filters of kernel size 15 and then transformed by a sigmoidal nonlinearity in the first layer. To implement the 15×15 linear filter, we stacked a sequence of linear convolutional layers with small filters (kernel size 3) in place of a single large convolutional layer, which we term the linearly-stacked convolutional layer [6]. This novel implementation of the convolutional layer outperforms the traditional manner of fitting each region in a large filter independently. One explanation is that this structure reduces the number of parameters, and also encourages a more localized filter which matches the tendency of retinal neurons to have a localized center-surround structure.

**Fig. 1:**
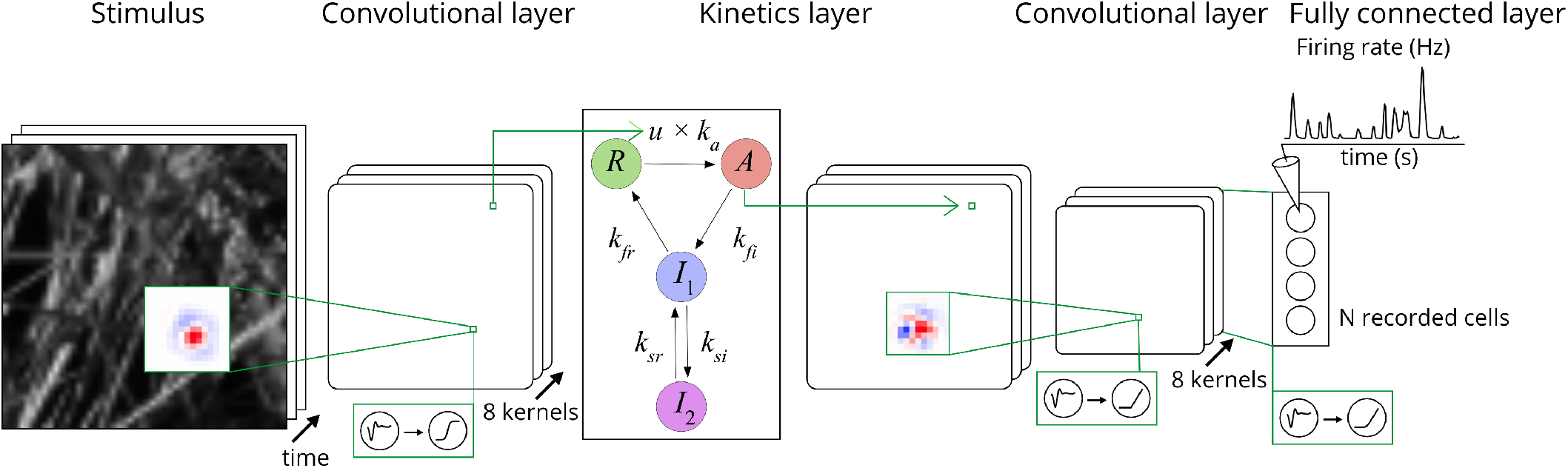
Model architecture. The model is based roughly on the structure of the retina in terms of three layers and the number of cell types in each layer, although there are more cell types in the retina than kernel types, and so the model is a simplification compared to the retina. The kinetics layer is placed at the location of the output synapses of the first layer, which would be roughly equivalent to bipolar cell synapses. Details are explained in the main text.

The second layer consists of recurrent synaptic units taken from the LNK model [1] that are capable of capturing slow dynamics of up to ten seconds. Each entry of the first layer output tensor serves as the input *u* of its own kinetic system while the transition rate parameters are shared across all entries. Each kinetic system contains four kinetic states where we denote *R, A, I*_1_, *I*_2_ the occupancies of the resting state, the active state, and the two inactivated states, respectively. The dynamics of each kinetic system is governed by the following master equation:

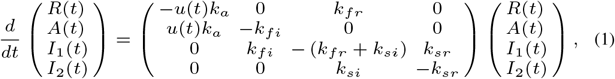

where *k*_*a*_, *k*_*fr*_, *k*_*fi*_, *k*_*si*_, and *k*_*sr*_ are kinetic parameters. The output of this kinetic layer is the tensor of active states occupancies A(*t*) whose shape is the same as that of the first layer.

The third layer consists of a linearly-stacked convolution with kernel size 11 and a rectifying nonlinearity in eight output channels. The final layer is a fully connected linear layer with a softplus activation function.

### E. Model training

As shown in Fig. 2, to fit slow and fast dynamics together, we first optimized all fast spatiotemporal parameters (all parameters except *k*_*si*_ and *k*_*sr*_), then separately optimized recurrent slow synaptic parameters (*k*_*si*_ and *k*_*sr*_).

**Fig. 2:**
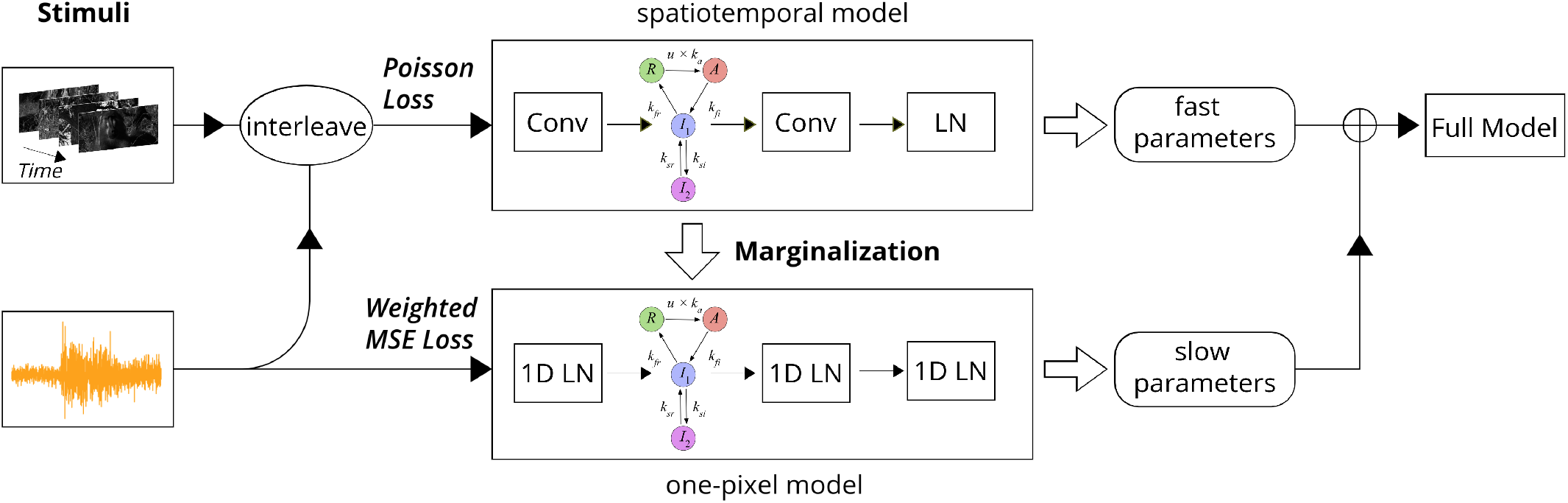
Schematic of training procedure. We first fit the fast spatiotemporal parameters using interleaved data, then fit the slow parameters in a marginalized one-pixel model using spatially uniform data. Details are explained in the main text.

First, we initialized the fast kinetic parameters (*k*_*a*_, *k*_*fr*_, *k*_*fi*_) according to the averaged values in the LNK model reported in [1]. We then interleaved the natural scene and the spatially uniform white noise datasets in the training dataset and used it to optimize all fast parameters with the truncated backpropagation through time (TBPTT) algorithm to minimize the Poisson loss function.

For fitting slow dynamics with the white noise dataset, we first marginalized the model into a one-pixel model given that the stimulus and the activation in every layer are spatially uniform. Specifically, the convolutional layers and the fully connected layer were marginalized into one-dimensional linear functions while the nonlinearities and the kinetic layer remained unchanged. We then froze all fast parameters and only optimized the slow parameters in the one-pixel model with the TBPTT algorithm. The marginalization reduced the model size dramatically and thus allowed the long truncation length (10 seconds in the data), which was essential for fitting slow dynamics.

To avoid the high-frequency component dominating the loss function when training the one-pixel model, we normalized the loss in different time bins and frequency components. We first divided the neural response *y*(*t*) and the model prediction *y*′(*t*) into 10 s time bins. Each segment was then filtered by a low pass and a high pass filter that were separated by a cutoff frequency dividing the response into two bins with approximately equal power. The loss function was then computed as

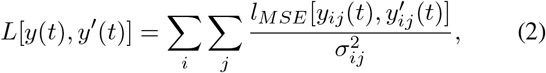

where *y*_*ij*_(*t*) 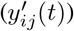 is the response (model output) for time bin *i* and frequency bin *j, σ*_*ij*_ is the standard deviation of *y*_*ij*_(*t*), *l*_*MSE*_(·,·) is the mean square error function.

Optimization was performed using ADAM [11] via Pytorch [12] on NVIDIA Titan X GPUs. The network was regularized with an L2 weight penalty at each layer.

## III. Results

Adaptive properties of model responses have been quantified by observing changes in the parameters of a simpler model fit under different stimuli, e.g. a linear-nonlinear (LN) model consisting of a linear temporal filter and a static nonlinearity, which is fit by the standard method of reverse correlation to the spatially uniform Gaussian white noise [4] (Fig. 3a). Note that the filter in this simple LN model are different from filters in the more complex model. Our model captures the fast change in temporal processing between low and high contrasts where the linear filter of an LN model at high contrast is faster and more biphasic than that of low contrast. Our model also captures the fast change in sensitivity as defined as the mean slope of the LN model nonlinearity as well as the fast and slow changes in offset as defined as the mean value of the nonlinearity. Furthermore, our model captures the multiscale contrast adaptation across a population of model units (Fig. 3c). In summary, the measure based on the LN model implies that our model accurately captures contrast adaptation over multiple timescales.

**Fig. 3:**
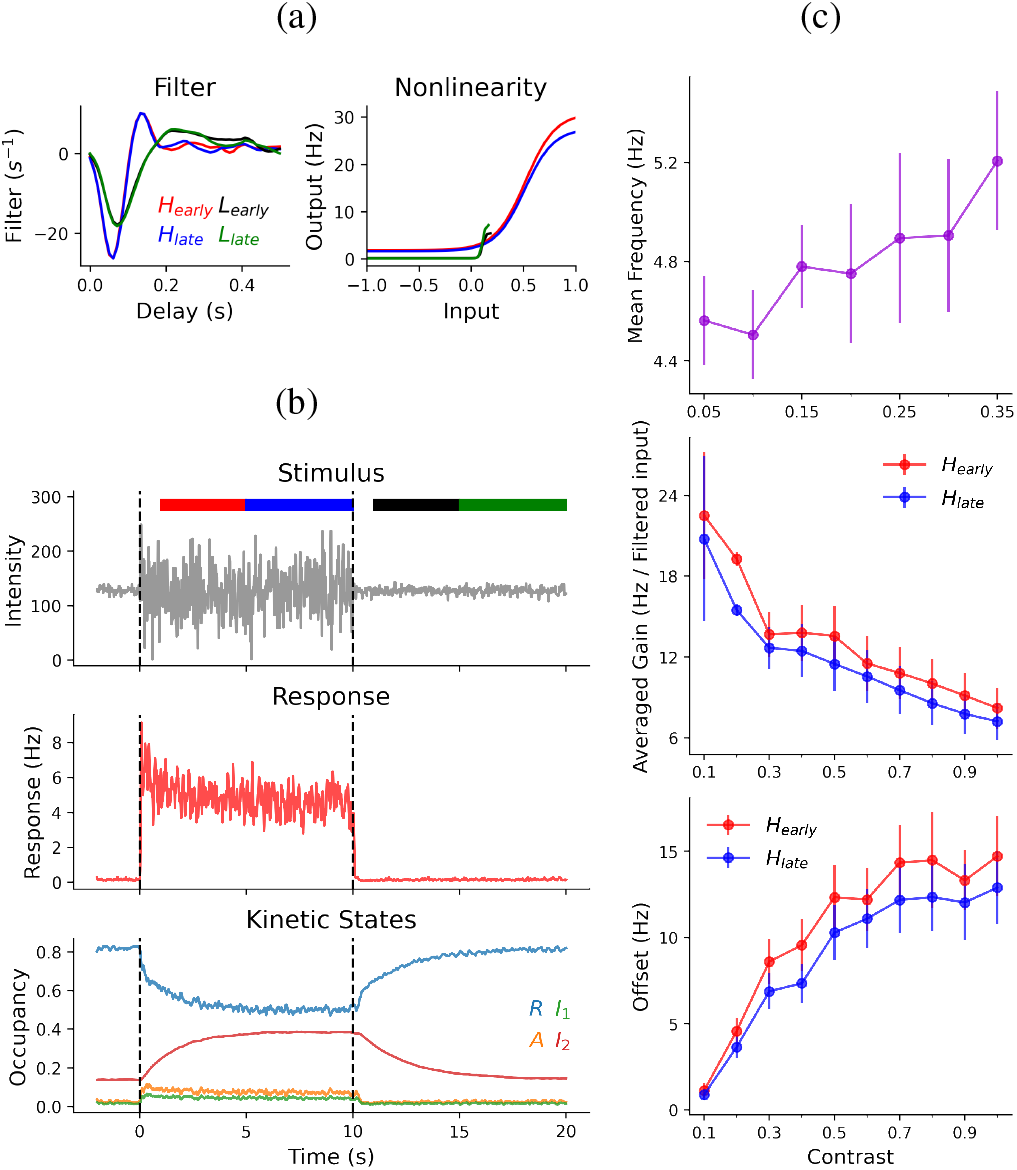
The model captures multiscale contrast adaptation. (a) LN models fit to the model output of different intervals indicated by the colored bars in (b) where low contrast is 0.05 and high contrast is 0.35. Left, linear temporal filters. Right, static nonlinearities. (b) Top, an example single trial of the spatially uniform Gaussian white noise stimulus. Colored bars indicate intervals *H*_*early*_, 1-5 s after the transition to high contrast, *H*_*late*_, 5 - 10 s after the transition to high contrast, *L*_*early*_ and *L*_*late*_, defined as similar time intervals after a low contrast step. Middle, the model response of one output unit averaged over 100 trials. Bottom, the occupancies of the kinetic states averaged over 100 trials for a contributing channel. The dashed lines indicate the transitions between stimuli of different contrasts. (c) Population summary of contrast adaptation in the model. Top, the mean frequency of the linear filter in LN models versus the contrast of the stimulus. Middle, the average slope of the nonlinearity in the LN models fit to *H*_*early*_ and *H*_*late*_ for different high contrasts while low contrast was held at 0.05. Bottom, average value of the nonlinearity in LN models fit to *H*_*early*_ and *H*_*late*_ for different high contrasts while low contrast was held at 0.05

Inspecting the occupancies of the kinetic states reveals how our model captures multiscale contrast adaptation (Fig. 3b). Overall, the first three states (*R, A*, and *I*_1_) reach equilibrium very quickly while the *I*_2_ state reaches equilibrium slowly, serving as a reservoir to control the total occupancy of the other three states. The gain of the kinetics layer, which is proportional to the resting state occupancy [1] drops rapidly after the switch from low contrast to high contrast because the mean value of the outgoing transition rate from the *R* state is increased by the high contrast stimulus. At the same time, the active state occupancy increases rapidly resulting in the strong model response just after the transition to high contrast. The *I*_1_ state occupancy also increases when reaching the equilibrium among the three fast states, leading to a gradual increase of the *I*_2_ state occupancy. As a consequence, the rising *I*_2_ state reduces gradually the total occupancy of the other three fast states producing the slow exponential decay of the model response at high contrast. For the switch from high contrast to low contrast, the mechanism is similar.

Our model also captures immediate spatiotemporal light responses. The model can predict spiking responses of salamander retinal ganglion cells accurately, approaching retinal reliability for both natural scenes and spatially uniform Gaussian white noise (Fig. 4). Moreover, the model performances drop significantly on shuffled datasets that destroy the temporal structure of the data longer than 0.4 seconds. This procedure confirms the recurrent nature of the model and implies that the recurrent structure in the model is functioning to accurately process information over longer timescales.

**Fig. 4:**
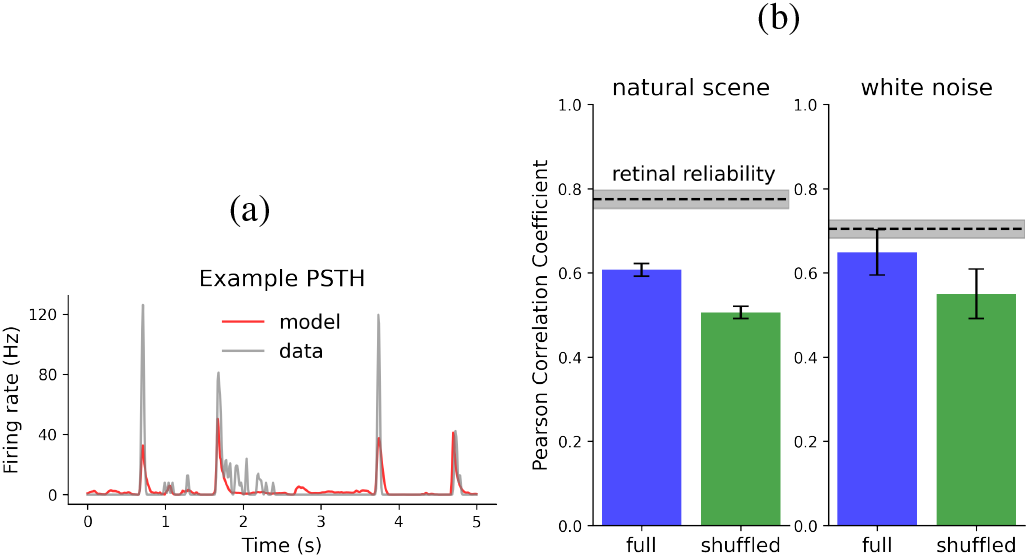
The model captures immediate light responses for both stimuli. (a) An example section of the Peristimulus time histogram (PSTH) of the model response and the recorded response. (b) The Pearson correlation measure of the model performance on both the natural scene and the spatially uniform Gaussian white noise. The performances on the normally ordered dataset and the randomly shuffled dataset are compared.

## IV. Conclusions

We created a mechanistically interpretable model that is capable of capturing contrast adaptation over multiple timescales as well as spatiotemporal immediate light responses to natural scenes. To optimize this model, we designed a novel training strategy for capturing fast and slow neural dynamics separately with a single model, which is difficult because hidden variables evolving over different timescales are entangled with each other, interact with the constantly fluctuating stimulus and are further nonlinearly combined to produce the observable output response.

Our results suggest a general approach to fitting multiscale dynamics that might be applicable to capturing other long-timescale phenomena, such as sensitization [10], [13] which has similar dynamics to contrast adaptation but opposite in sign, where certain cells become more sensitive when stimulated with a localized strong stimulus. Compared to previous CNN models that only capture short-term neural dynamics for retinal ganglion cells, the additional kinetics blocks in our model act as recurrent structures to capture long-term dynamics. This kinetic structure has a strong correspondence to the biophysics of retinal synapses, following the general principle of constraining the model architecture to be more realistic through the demand of replicating neural phenomena.

## Acknowledgments

The authors would like to thank Joshua Melander, Michael Menz and our lab members for helpful discussions and experimental assistance, as well as Surya Ganguli for helpful discussions. This work was supported by grants R01EY022933, R01EY025087 and P30EY026877 to SAB from the NIH (NEI). We would also like to thank the Methods in Computational Neuroscience course at the Marine Biological Laboratory.

